# Distributed and Retinotopically Asymmetric Processing of Coherent Motion in Mouse Visual Cortex

**DOI:** 10.1101/791905

**Authors:** Kevin K. Sit, Michael J. Goard

**Affiliations:** Department of Psychological & Brain Sciences, University of California, Santa Barbara, Santa Barbara, CA 93106, USA; Department of Molecular, Cellular, and Developmental Biology, University of California, Santa Barbara, Santa Barbara, CA 93106, USA; Neuroscience Research Institute, University of California, Santa Barbara, Santa Barbara, CA 93106, USA

**Keywords:** Motion perception, mouse vision, higher visual areas, calcium imaging

## Abstract

Perception of visual motion is important for a range of ethological behaviors in mammals. In primates, specific higher visual cortical regions are specialized for processing of coherent visual motion. However, the distribution of motion processing among visual cortical areas in mice is unclear, despite the powerful genetic tools available for measuring population neural activity. Here, we used widefield and 2-photon calcium imaging of transgenic mice expressing a calcium indicator in excitatory neurons to measure mesoscale and cellular responses to coherent motion across the visual cortex. Imaging of primary visual cortex (V1) and several higher visual areas (HVAs) during presentation of natural movies and random dot kinematograms (RDKs) revealed heterogeneous responses to coherent motion. Although coherent motion responses were observed throughout visual cortex, particular HVAs in the putative dorsal stream (PM, AL, AM) exhibited stronger responses than ventral stream areas (LM and LI). Moreover, beyond the differences between visual areas, there was considerable heterogeneity within each visual area. Individual visual areas exhibited an asymmetry across the vertical retinotopic axis (visual elevation), such that neurons representing the inferior visual field exhibited greater responses to coherent motion. These results indicate that processing of visual motion in mouse cortex is distributed unevenly across visual areas and exhibits a spatial bias within areas, potentially to support processing of optic flow during spatial navigation.

## INTRODUCTION

Perception of visual motion is critical for animal survival, underlying behaviors such as visually guided navigation, pursuit of prey, and avoidance of threats. Although neurons selective for visual motion arise early in the visual system, extensive research in primates has shown that perception of coherent global motion independent of local motion relies on processing in specialized regions of visual cortex [1,2]. The cortical processing of coherent motion has been studied extensively in primates, but is not as well understood in the mouse, [1,2] although mouse visual cortical neurons are known to be well-tuned for coherent visual motion [3,4], and vision plays an important role in navigation [5], but the cortical organization of coherent motion processing is poorly understood. Recently developed techniques for measuring and manipulating neural activity in genetically identified neurons makes the mouse an attractive model system for investigating the neural circuitry underlying coherent motion processing [6,7]. Although visually-driven neurons along the retinogeniculocortical pathway of mice and primates exhibit differences in response properties and connectivity [8], there are parallels in the overarching meso- and macroscale organization of the visual areas [6]. Researchers are just beginning to understand how coherent motion is encoded in the mouse visual system, and the degree to which circuits underlying motion processing are homologous between mice and primates.

In primates, the middle temporal area (MT) and the downstream medial superior temporal area (MST) have been identified as specialized regions for processing of coherent motion [9], and serve as the gateway to the dorsal stream of visual processing [10,11]. Area MT contains a high proportion of direction selective neurons [12–14] and preferentially receives direction selective inputs from the primary visual cortex (V1) [15]. Pharmacological lesioning of MT causes deficits in coherent motion perception [16], and microstimulation of MT can influence perception of motion using random dot kinematograms (RDKs) [17]. Individual neurons in MT and MST exhibit strong direction-selective responses to RDKs and many are selective for the overall motion of plaid stimuli (pattern direction-selective) rather than to the individual component gratings (component direction-selective) [9]. In contrast, neurons in the primate V1 are mostly nonselective for the direction of coherent motion in RDKs [18] and exclusively exhibit component direction-selective responses to plaids [9]. These findings have led to models in which MT response properties derive from weighted summation and normalization of direction-selective V1 inputs [19,20].

The basic organization of mouse visual cortex is similar to the primate visual system, with the majority of cortical input arriving via the retinogeniculocortical pathway (along with indirect input from the superior colliculus via thalamic nuclei [21]), and a network of hierarchically organized visual cortical regions with independent retinotopic maps [22–24]. However, the mouse visual system has several functional properties that are distinct from that of primates. For example, several types of mouse retinal ganglion cells (RGCs) exhibit direction selectivity [25–27], while thus far direction-selective RGCs have not been found in primate retina [28]. Direction-selective RGCs have an asymmetric retinotopic distribution [29,30], give rise to direction-selective inputs to visual cortex [31], and influence direction-selectivity in visual cortex [32]. In addition, strong orientation tuning and direction selectivity are already present in the lateral geniculate nucleus [33–35], in contrast to the weakly tuned LGN neurons found in primates [36]. Finally, in contrast to primate V1 [9,18], mouse V1 contains a significant fraction of neurons that exhibit tuned responses to global coherent motion found in RDKs [3] and plaid pattern motion ([37,38], though see [39]).

In recent years, mapping procedures using intrinsic signal imaging [23,24,40] and widefield calcium imaging [41,42] have allowed researchers to define and functionally characterize higher visual areas (HVAs) in intact mice, but the functional role of the HVAs, and any homology to primate visual structures, remains an area of active investigation. Anatomical and functional studies have found that HVAs are broadly connected into two subnetworks with projection patterns similar to primate ventral and dorsal streams [40,43–45]. Specifically, the lateral medial (LM) and lateral intermediate (LI) areas preferentially project to temporal and lateral entorhinal cortices (putative ventral stream) while the anterolateral (AL), posterior medial (PM), rostrolateral (RL), and anteromedial (AM) areas preferentially project to parietal, motor, and medial entorhinal cortices (putative dorsal stream). Consistent with this classification, measurements of single neuron activity indicated greater direction selectivity in regions AL, RL, and AM ([23], though see [24]), a hallmark of dorsal stream regions in primate [12–14].

Here, we used widefield and 2-photon calcium imaging to map areal and cellular responses to coherent motion in mouse visual cortices using both natural movies and RDKs. We found that HVAs exhibit heterogeneous responses to coherent motion as in primates, with stronger activation in response to coherent motion in regions AL, PM, and AM compared to V1, LM, and LI. On the other hand, responses to coherent motion were much more distributed than in primate, with neurons in all measured regions (including V1) containing a significant fraction of coherent motion responsive cells. Furthermore, coherent motion responses were distributed asymmetrically across visual elevation, both within and across all visual regions, with neurons representing the inferior visual field exhibiting much stronger coherent motion responses. Taken together, these results show that the mouse visual cortex is optimized for distributed processing of inferior field motion, potentially enhancing processing of optical flow signals during movement.

## RESULTS

### Mouse visual cortex exhibits heterogeneously distributed responses to coherent motion

In order to measure neural responses to coherent motion in an unbiased manner across the visual cortex, we used a custom widefield microscope (Figure 1A Methods) to measure calcium responses through a 4mm diameter window located over the visual cortex of awake, head-restrained mice expressing the calcium indicator GCaMP6s in excitatory neurons (Emx1-Cre::Rosa-tTA::TITL-GCaMP6s [46,47]; see Methods; Supplemental Video S1). Using established mapping procedures for defining visual HVAs [23,40] that were adapted for calcium imaging [41,42] (see Methods), we determined the areal boundaries of primary visual cortex (V1) and six consistently identified higher visual areas (HVAs): LM, AL, PM, LI, RL, and AM (Figure 1B), ordered by their approximate position in the visual hierarchy [44]. As in some previous studies, we were not consistently able to locate area A independent of AM [40], so this area was not included in our analyses. This procedure was performed separately for each mouse to obtain a precise map of the visual cortical areas (Figure S1). To determine which areas responded to complex visual input, we played repeated presentations of sets of natural movies recorded from a head-mounted camera [48] (Figure 1C) on a large monitor subtending 130° azimuth (0 to 130° nasal to temporal) and 100° elevation (−50° to +50° inferior to superior) of the contralateral visual field. Calcium responses from individual pixels in visual cortex exhibited reliable responses to repeated presentations of the natural movies (Figure 1D). To reduce inter-mouse variability and hemodynamic artifacts from blood vessels, we aligned HVA area boundaries and averaged reliability across multiple mice (Figure 1E; *n* = 19 sessions across 7 mice), revealing that primary visual cortex exhibited uniformly reliable responses across the region. Response reliability was slightly weaker in secondary visual regions LM and PM, and weaker still in higher areas of the visual hierarchy such as AL, RL, and AM (Figure S2).

**Figure 1:**
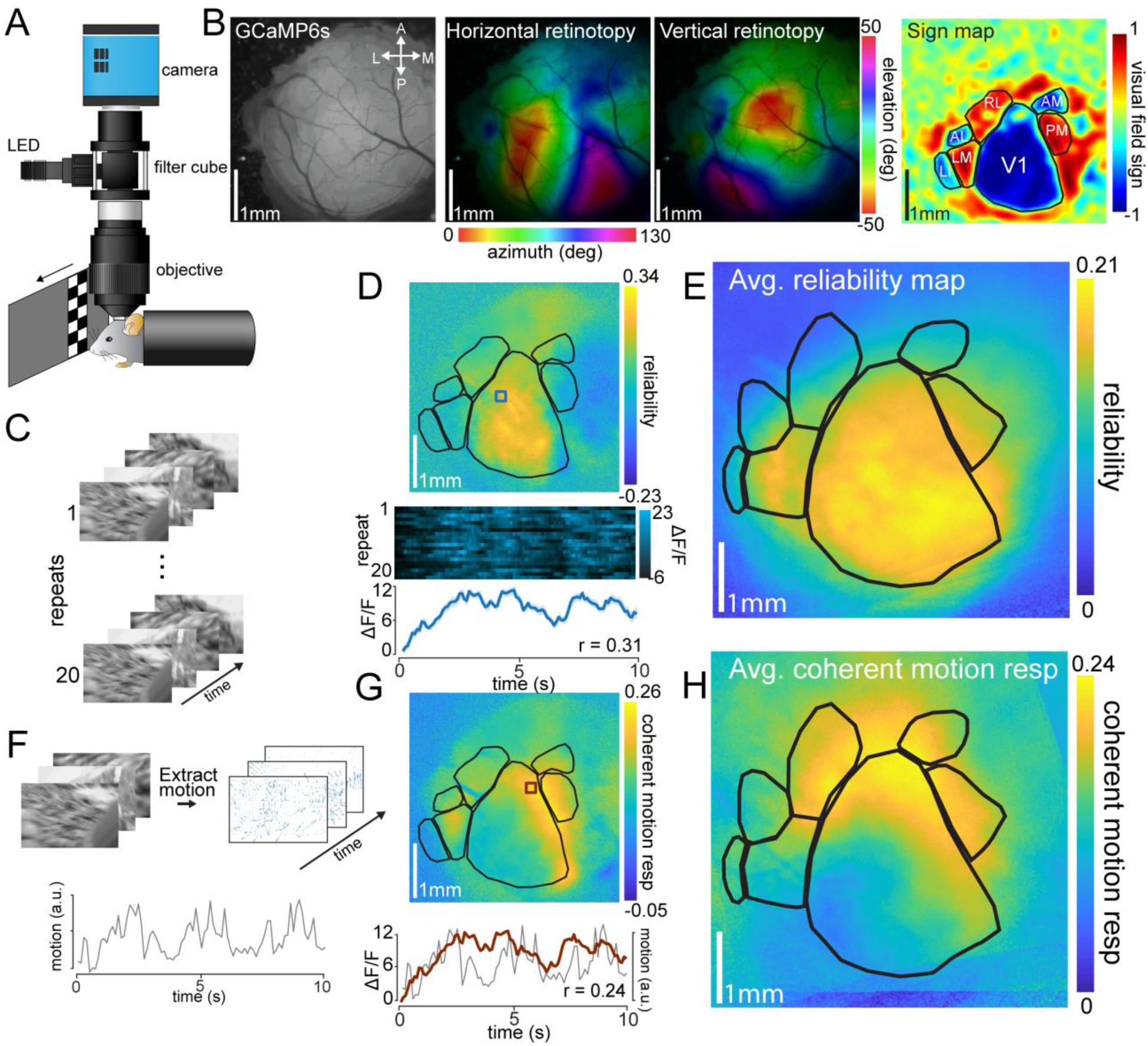
Mesoscale calcium responses to coherent motion in natural visual stimuli. **(A)** Schematic of the custom epifluorescent widefield microscope for *in vivo* GCaMP6s imaging. The screen depicts a retinotopic mapping stimulus, with a drifting bar moving across the visual field (azimuth mapping). **(B)** Areal maps from an example mouse. Left: surface raw fluorescence image of a 4mm cortical window of example Emx1-GCaMP6s mouse. Middle: horizontal and vertical retinotopic maps showing preferred location of each pixel for azimuth (left) and elevation (right); color bars indicate degree offset from center of visual field). Imaging plane is 500µm ventral from the surface of cortex. Right: sign map (red, positive; blue, negative), and resulting segmentation of visual cortex into V1 and HVAs. Boundaries are calculated using the reversal of visual field sign and nonredundant coverage of visual space (see Methods). Scale bars = 1 mm. **(C)** Schematic of the natural movie stimulus. Scenes were repeated 20 times to measure reliable neural responses. **(D)** Top: map from a single mouse showing reliability across posterior cortex; visual area segmentation as in (B). Bottom: multi-trial response (20 repeats) and mean trace (± s.e.m. shaded) of a single pixel (blue square) to repeated presentation of the natural movie. Reliability is defined as the between-trial correlation coefficient (*r* = 0.31). **(E)** Mean reliability map across all imaged mice (*n* = 19 sessions over 7 mice). Individual maps are transformed onto a common coordinate system for comparison across mice. **(F)** Extraction of the coherent motion in the stimulus. Pixel-wise motion vectors were extracted from each frame of the movie, and the sum of these vectors is used as a measure of the net coherent motion of each frame. **(G)** Top: map from a single mouse showing motion response across posterior cortex; visual area segmentation as in (B). Bottom: Neural response of a single pixel (red) overlaid on the motion trace (gray); pixel-wise motion response is calculated as the Pearson correlation between these two signals (*r* = 0.24). **(H)** Mean motion response map across all imaged mice (*n* = 19 sessions over 7 mice); alignment procedure same as (E). Area abbreviations: primary visual (V1), lateral medial (LM), anterolateral (AL), posterior medial (PM), laterointermediate (LI), rostrolateral (RL), and anteromedial (AM).

We next investigated whether the responses to coherent motion embedded within the natural movies were similarly uniform across areas, or if the mouse exhibits heterogeneity across HVAs as has been observed in primate MT/MST. To this end, we first calculated the total coherent motion for each frame of the natural movie (Figure 1F; Methods). We next measured the Pearson correlation between the deconvolved neural response for each pixel and the coherent motion energy of the scene (Figure 1G, bottom; Methods). In both individual sessions (Figure 1G, top) and aligned session averages (Figure 1H; *n* = 19 sessions across 7 mice), we found that particular cortical locations were strongly driven by coherent motion embedded in the scene. The motion response map (Figure 1H) was not well correlated to the map of reliability to all visual features (Figure 1E), as evident by the low pixel-wise correlation between motion response and reliability (Figure S3; *r* = 0.088 ± 0.12, *p* = 0.47, mean ± s.e.m., single sample t-test).

One possibility for the heterogeneous distribution of motion responses in visual cortex is that motion energy is not uniformly distributed in the natural movies. Indeed, individual natural movies exhibit non-uniform distributions of motion energy (Figure S4), though the average motion energy across all natural movies is roughly uniform (Figure S4). To further ensure that the distribution of motion energy is unbiased across the visual field, and to remove the influence of other features driving neural responses, we used full screen random dot kinematograms (RDKs) to measure responses to coherent motion across the visual field (Figure 2A; Methods).

**Figure 2:**
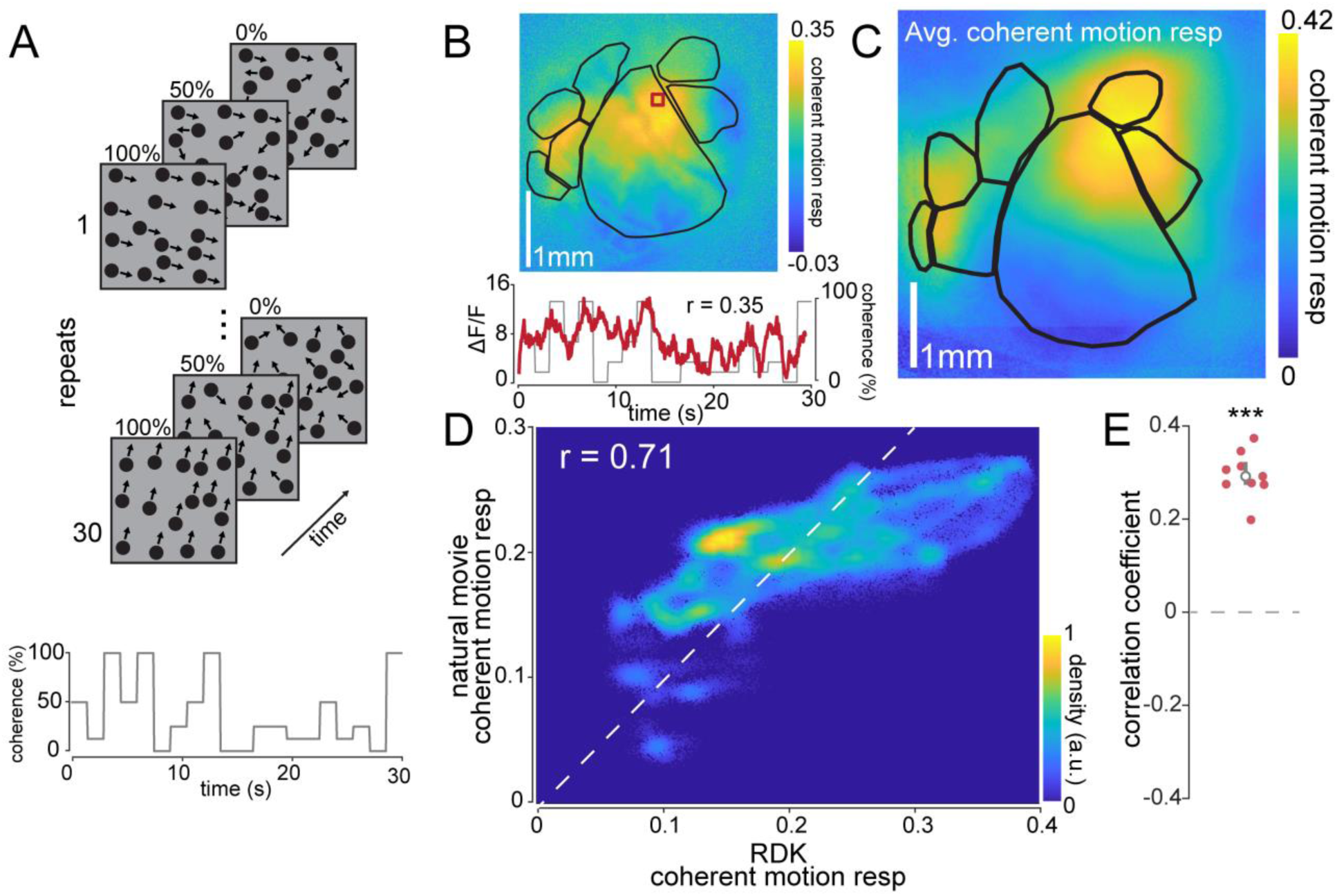
Mesoscale calcium responses to coherent motion in RDKs. **(A)** Schematic of the RDK stimulus. A single repeat of the stimulus contains dots drifting at varying coherence values across the trial duration (bottom). Over multiple repeats, the coherence values remain constant while the dot positions and directions are randomized every trial. **(B)** Top: map from a single mouse showing motion response to RDKs across posterior cortex. Bottom: Single pixel response (red) over the course of a single presentation of the stimulus overlaid on the coherence value trace; pixel-wise motion response is calculated as the Pearson correlation between these two signals (*r* = 0.35). **(C)** Mean motion response map across all imaged mice (*n* = 10 sessions over 10 mice). Individual maps are transformed onto a common coordinate system for comparison across mice. **(D)** Pixel-wise density plot of RDK vs. natural movie motion response across all mice to show correlation of motion response across visual stimuli (*r* = 0.71). Density plots are similar to scatter plots, but provide additional information about the density (but not strength) at each coordinate, with warmer colors indicating a higher density of observation pairs (see Methods). **(E)** Natural movie versus RDK motion response correlation for each mouse (*r* = 0.30 ± 0.02, *n* = 10, *p* = 1.0 × 10^−7^).

The calcium response within a single pixel represents the summed activity across many neurons, so we could not measure tuning to preferred motion direction as is typically done in single neuron recordings. Instead, we took advantage of the fact that RDKs with higher coherence values will strongly drive neurons responsive to coherent motion and result in higher magnitude calcium responses regardless of motion direction. We used a small dot size (2° of visual space) and randomized dot position and drift direction on each trial, so that the only consistent visual feature across trials was the RDK motion coherence. We then defined the *coherent motion response* for each pixel as the Pearson correlation coefficient between the pixel response (ΔF/F) and the RDK motion coherence (see Methods). We found that individual pixels in V1 and other HVAs exhibited a high degree of correlation with RDK coherence (Figure 2B). Averaging and aligning motion response maps revealed a heterogeneous distribution of coherent motion responsive pixels within and across regions (Figure 2C). Despite the difference in stimulus, the motion response map measured with RDKs (Figure 2C) was well correlated to the motion response map measured with natural movies (Figure 1G), as indicated by the Pearson correlation of aligned pixels (Figure 2D-E; *r* = 0.71; *p* = 1.0 × 10^−7^, single sample t-test), although certain regions (e.g., RL) showed differences between stimuli (see Discussion).

### Higher visual areas exhibit differential responses to coherent visual motion

In primates, particular higher visual areas (MT/V5 and MST) are specialized for processing of coherent motion. To determine if particular mouse HVAs exhibit a similar degree of dedicated motion processing, we examined coherent motion responses in V1, each consistently defined HVA (LM, PM, AL, LI, RL, and AM), and somatosensory region S1 as a negative control.

For each session, we used retinotopic mapping to define visual areas, and then measured coherent motion responses for each pixel as the Pearson correlation between the deconvolved ΔF/F signal and the RDK motion coherence (Figure 3A, B). We then averaged all of the pixels within each defined region of interest to generate a measure of coherent motion response by area, revealing that all visual regions except LI exhibited coherent motion response (R_CM_) values significantly above zero (Figure 3C, V1: R_CM_ = 0.08 ± 0.02, *p* = 3.0 × 10^−3^; LM: R_CM_ = 0.09 ± 0.03, *p* = 9.7 × 10^−3^; AL: R_CM_ = 0.17 ± 0.03, *p* = 1.1 × 10^−4^; PM: R_CM_ = 0.15 ± 0.03, *p* = 3.0 × 10^−4^; LI: R_CM_ = 0.03 ± 0.02, *p* = 0.11; RL: R_CM_ = 0.11 ± 0.02, *p* = 1.1 × 10^−3^; AM: R_CM_ = 0.21 ± 0.03, *p* = 1.0 × 10^−4^; mean ± s.e.m., single-sample t-test). As expected, area S1 was not motion responsive (R_CM_ = 0.003 ± 0.01, *p* = 0.59; mean ± s.e.m., single-sample t-test). Moreover, some HVAs had significantly higher coherent motion response than other visual regions (Figure 3D). Specifically, area AM had significantly higher motion response values than all other regions, followed by AL, PM, RL, V1, and LM, while area LI had the lowest coherent motion response (schematized in Figure 3E). These results indicate that particular HVAs in mice indeed exhibit enhanced responses to coherent motion relative to primary visual cortex. Given the differences in coherent motion responses in HVAs, these results are broadly consistent with past anatomical and functional work suggesting that areas LM and LI constitute a homologue of the ventral stream, while areas AL, PM, RL, and AM constitute a homologue of the dorsal stream [43–45].

**Figure 3:**
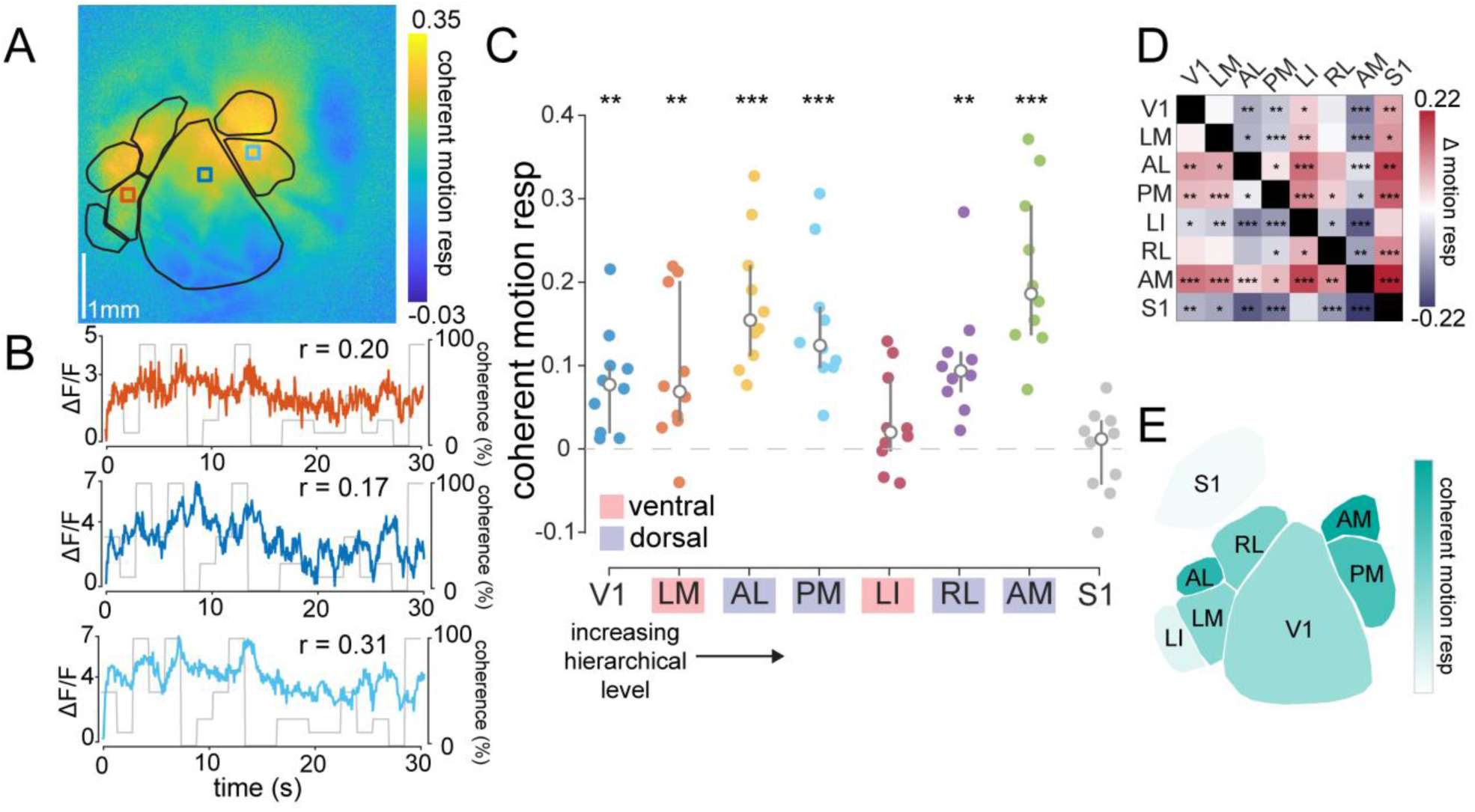
HVAs exhibit heterogeneous coherent motion responses to RDKs. **(A)** Map from a single mouse highlighting heterogeneous coherent motion responses across HVAs. **(B)** Single pixel responses taken from area LM (orange; top), V1 (dark blue; middle), and PM (light blue; bottom) overlaid on the coherence trace. Correlation between pixel response and coherent motion trace indicated for each pixel. **(C)** Across all mice (*n* = 10 sessions over 10 mice), each tested visual area is significantly motion responsive, but a control area outside of visual cortex (region S1) is not (\**p* < 0.05, \*\**p* < 0.01, \*\*\**p* < 0.001; single-sample t-test). Overlay indicates median ± quartiles. **(D)** Relative difference matrix, with each element equal to the coherent motion response in the column region subtracted from the coherent motion response in the row region across mice (red, higher; blue, lower; \**p* < 0.05, \*\**p* < 0.01, \*\*\**p* < 0.001; two-sample paired t-test). **(E)** Simplified schematic of mean coherent motion responses in each cortical area across mice.

### Coherent motion responses exhibit retinotopic asymmetry

In addition to differences in coherent motion responses between visual areas, there was also heterogeneity within individual visual areas. To quantify this, we first looked at area V1, where pixels located in anterior V1 were more strongly driven by coherent motion than pixels in posterior V1 (Figure 4A, B). The heterogeneity of coherent motion responses in V1 could be explained in two ways: an anatomy-based organization (along anatomical axes) or a functional-based organization (along retinotopic axes). To test these possibilities, we isolated the entire visual cortex and calculated the z-scored motion response for each pixel as a function of azimuth and elevation, as mapped with drifting bar stimuli (Figure 4C). Although there was little correlation across mice (*n =* 10 sessions over 10 mice) between the z-scored motion response and the azimuth, there was a strong negative correlation between the motion response and the elevation, indicating stronger coherent motion processing in the inferior visual field. (Figure 4D-F; azimuth *r* = 0.09 ± 0.09, *p =* 0.34; elevation *r* = −0.54 ± 0.06, *p* = 1.91 × 10^−5^; single sample t-test)

**Figure 4:**
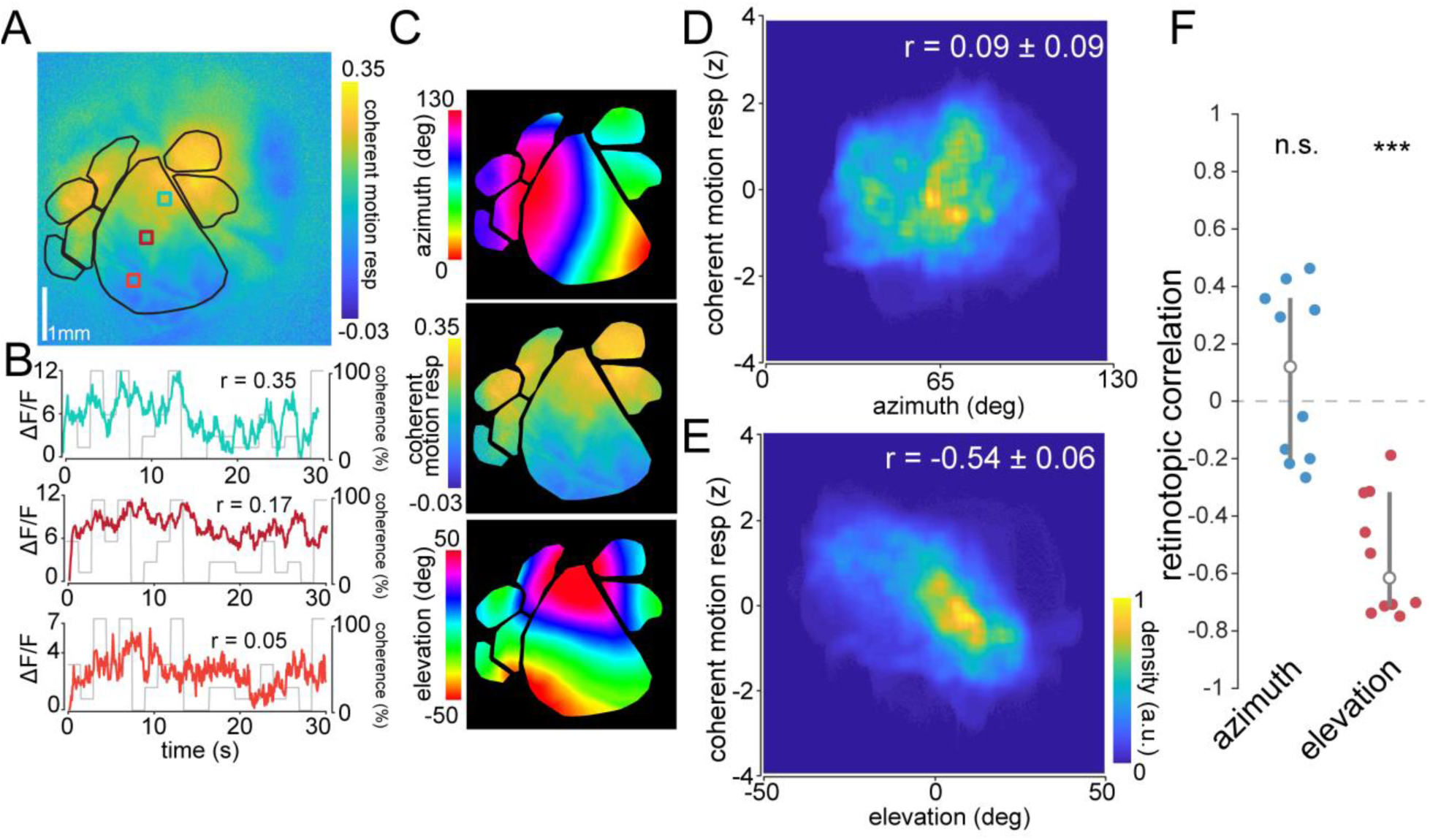
Retinotopic asymmetry of coherent motion responses across the visual cortex. **(A)** Map from single mouse showing gradient of coherent motion response along the vertical retinotopic axis of V1. **(B)** Single pixel responses from anterior (blue, *r* = 0.35), middle (red, *r* = 0.17), and posterior (blueorange *r* = 0.05) pixels in V1, showing different amounts of coherent motion response. **(C)** Visual cortex isolated maps from mouse in (A) of azimuth (top) and elevation (bottom), as well as the coherent motion response (middle) for comparison. **(D)** Density scatter plot across all mice (*n* = 10 sessions over 10 mice), showing no significant correlation to azimuth (*r* = 0.09 ± 0.09). Coherent motion response is z-scored to account for differences between experiments. **(E)** Same as (D), but for elevation, showing a significant negative correlation between elevation preference and coherent motion response across visual cortex (*r* = −0.54 ± 0.06). **(F)** Comparison of retinotopic correlation between azimuth and elevation. There is a significant anticorrelation of coherent motion response and elevation across mice, but none with azimuth. Error bars are mean ± quartiles (***: *p* < 0.001).

We next investigated whether the retinotopic asymmetry was also present in other HVAs. For each area, we calculated the correlation between the z-scored coherent motion response and azimuth (Figure 5A-H, left plots) and elevation (Figure 5A-H, right plots). Most of the areas exhibited no correlation between the preferred azimuth and the coherent motion response, with the exception of area LM and LI, which exhibited a significant positive correlation (Figure 5I; V1: *r =* 0.09 ± 0.12, *p* = 0.44; LM: *r =* 0.46 ± 0.13, *p* = 0.007; AL: *r* = −0.10 ± 0.14, *p* = 0.46; PM: *r* = −0.12 ± 0.13, *p =* 0.38; LI: *r* = 0.35 ± 0.10, *p =* 0.008; RL: *r* = −0.06 ± 0.20, *p =* 0.76; AM: *r* = −0.09 ± 0.12, *p =* 0.46; S1: *r* = 0.13 ± 0.13, *p =* 0.33; mean ± s.e.m., single-sample t-test). Conversely, almost all regions exhibited a significant negative correlation between the preferred elevation and the coherent motion response, with the exception of AL, as well as areas LI and S1—which both had little coherent motion response throughout the area (Figure 5J; V1: *r* = −0.81 ± 0.05, *p* = 1.4 × 10^−6^; LM: *r* = −0.44 ± 0.08, *p* = 4.3 × 10^−3^; AL: *r* = 0.03 ± 0.14, *p* = 0.85; PM: *r* = −0.78 ± 0.06, *p* = 4.5 × 10^−6^; LI: *r* = −0.04 ± 0.11, *p =* 0.73; RL: *r* = −0.27 ± 0.10, *p* = 0.035; AM: *r* = −0.36 ± 0.14, *p* = 0.036; S1: *r* = −0.14 ± 0.09, *p* = 0.11;mean ± s.e.m., single-sample t-test). Note that the negative correlation between the preferred elevation and motion response could not be explained by a decreasing anterior-to-posterior anatomical gradient, as several HVAs (e.g., area AM) had vertical retinotopic gradients oriented counter to the anatomical gradient. Furthermore, the negative correlations tended to be strongest in regions in which the pixels subtended a larger range of elevation (Figure S5), as measured by the elevation covered by 95% of the pixels (E_95%_), such as V1 (E_95%_ = 58.42 degrees) and PM (E_95%_ = 32.16 degrees), compared to areas AM (E_95%_ = 25.25 degrees) and AL (E_95%_ = 20.85 degrees). There was no relationship between azimuth correlations and azimuth coverage (A_95%_; Figure S5C). Although the retinotopic coverage of the HVAs is similar to that reported with intrinsic signal imaging in previous work [40], it is possible that the population averaging and low-pass filtering intrinsic to widefield population imaging approaches may underestimate the retinotopic extent of the individual neurons within each region.

**Figure 5:**
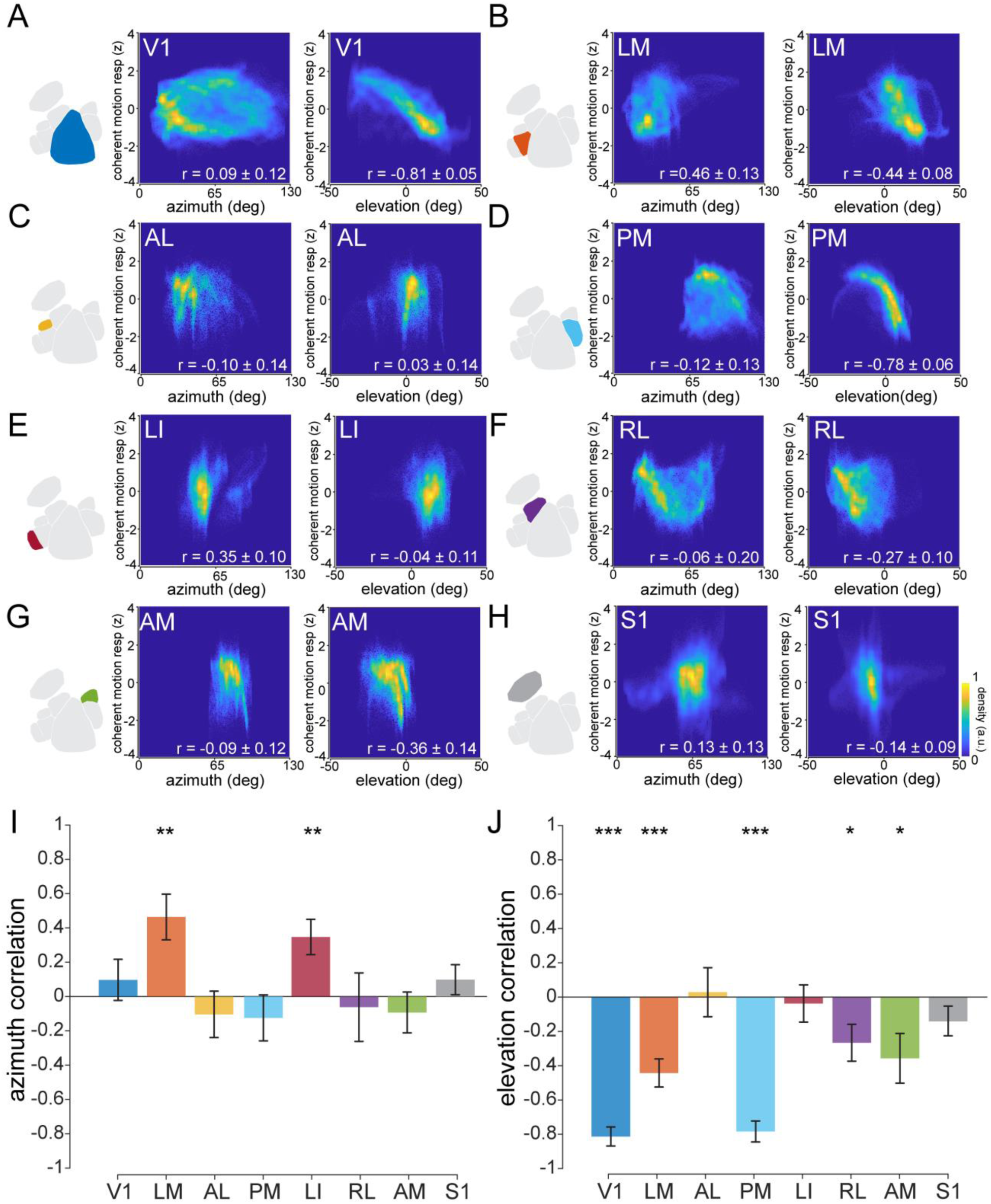
Retinotopic dependence of coherent motion response in each visual area. **(A)** Left: Combined density scatter plot across all mice (*n* = 10 sessions over 10 mice) comparing z-scored coherent motion response to azimuth (Pearson’s correlation coefficient *r* = 0.09, *p* = 0.44) for area V1. Right: same as left, but for elevation (*r* = −0.81, *p* < 0.001). **(B-H)** Same as (A) for each HVA (LM, AL, PM, LI, RL, AM), and somatosensory cortex (S1). **(I)** Mean Pearson’s correlation coefficient between azimuth and coherent motion response for each area; only area LM and LI have a significant correlation (LM: *r* = 0.46, *p* = 0.004; LI: *r* = 0.35, *p* = 0.008; for all other areas *p* > 0.37). **(J)** Same as (I), but for elevation. All visual areas except AL (*r* = 0.03, *p* = 0.85) and LI (*r* = −0.04, *p* = 0.74) show significant negative correlation between elevation and coherent motion response (*r* < −0.27; *p* < 0.05). Control area S1 does not exhibit a correlation for either azimuth (*r* = 0.13 *p* = 0.33) or elevation (*r* = −0.14, *p* = 0.14). Single sample t-test (\**p* < 0.05; \*\**p* < 0.01; \*\*\**p* < 0.001). Bars are mean ± s.e.m.

### Coherent motion responses of single neurons in mouse visual cortex

Although widefield imaging revealed differences in coherent motion responses both across and within visual regions, we wanted to confirm that individual neurons exhibited the same response distributions. To this end, we used 2-photon calcium imaging to measure the activity of populations of Layer 2/3 excitatory neurons in regions V1 and a subset of HVAs (LM, PM, and AM) that exhibited varying levels of coherent motion response (Figure 6A-B). To determine whether individual neurons exhibit coherent motion responses, we displayed RDKs in 8 directions at a range of coherence values (0 - 100%), then measured the correlation coefficient between the average calcium response and the coherence trace for each direction (Figure 6C; Supplemental Video S2). We found that individual neurons showed significant coherent motion selectivity in V1 (74% of 11,018 cells, *n* = 27 sessions over 15 mice) and all imaged HVAs, including LM (83% of 2166 cells, *n* = 8 sessions over 8 mice), PM (84% of 1781 cells, *n* = 8 sessions over 8 mice), and AM (82% of 1178 cells, *n* = 7 sessions over 7 mice). As expected, individual neurons exhibited selectivity for particular coherent motion directions [3] (Figure 6B-C). As such, across all visual areas, population responses to direction were highly mixed, though there was a slight bias toward the horizontal directions (nasal and temporal; Figure 6D-E) that was consistent across imaging fields and mice (Figure 6F; *p* = 1.76 × 10_^-6^_, *n* = 3,461 cells, Hodges-Ajne test for circular non-uniformity). Although the mean coherent motion response for the preferred direction was significant for all visual areas imaged (Figure 6G; V1: R_CM_ = 0.26 ± 0.02, *p* = 1.6 × 10^−17^; LM: R_CM_ = 0.31 ± 0.01, *p* = 1.3 × 10^−5^; PM: R_CM_ = 0.37, ± 0.01, *p* = 3.6 × 10^−6^; AM: R_CM_ = 0.36 ± 0.01, *p* = 2.2 × 10^−6^; mean ± s.e.m., single sample t-test), it was significantly higher in areas PM (*p* = 1.3 × 10^−2^, two sample t-test) and AM (*p* = 1.7 × 10^−3^, two sample t-test) as compared to V1, similar to results from widefield imaging (Figure 3).

**Figure 6:**
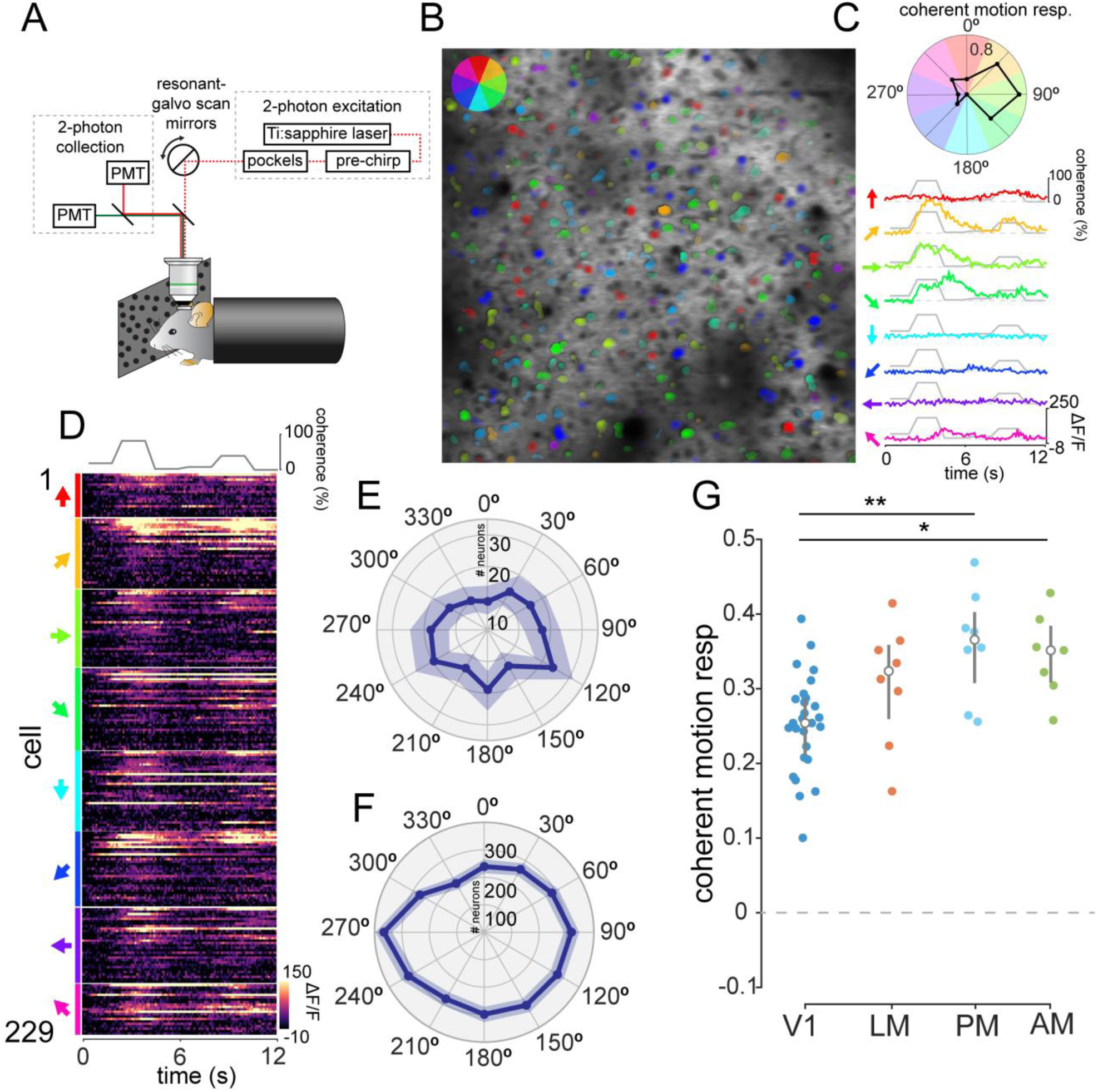
Coherent responses of single cells. **(A)** Schematic of the two-photon microscopy experimental setup. **(B)** Example two-photon imaging field from V1. Each neuron is color-coded by its preferred direction; the opacity of the color corresponds to its mean activity level. **(C)** Top: Polar plot of coherent motion response for each motion direction. Bottom: Responses of a single example neuron to each of the 8 motion directions, highlighting the direction selectivity of an individual neuron. The coherence trace for this session is overlaid on each neural response. **(D)** Responses of each cell in the example session, grouped by preferred direction (left, arrows) and ordered by decreasing coherent motion response. **(E)** Polar plot of averaged directional tuning to coherent motion across all cells in the example session. **(F)** Histogram of 30 degree binned preferred direction of all neurons across all sessions. Note the significant non-uniformity across all sessions (*n* = 3461, *p* = 1.73 × 10^−6^, Hodges-Ajne test for non-uniformity). **(G)** Mean coherent motion response across HVAs. Each point represents the mean coherent motion response of all the cells in that session. Although all areas exhibit significant coherent motion responses (*p* < 1.3 × 10^−5^), areas AM and PM have significantly higher coherent motion responses than area V1 (\**p* < 0.05; \*\**p* < 0.01; \*\*\**p* < 0.001, two sample t-test).

### Retinotopic asymmetry in coherent motion response also evident in individual neurons

Finally, we wanted to confirm that the inferior field bias of coherent motion processing evident in the widefield imaging experiments is also present at the level of individual neurons. To test this, we first measured the preferred azimuth and elevation of populations of V1 neurons using flashing bars with a flickering checkerboard pattern randomly presented at a range of locations (Figure 7A; see Methods). We then measured the coherent motion response in the preferred direction for each neuron (Figure 7B-C) and correlated the coherent motion response with the preferred azimuth and elevation. Similar to the widefield imaging results (Figure 4), neurons preferring lower elevation exhibited greater coherent motion responses than neurons preferring higher elevation (Figure 7D-E), although the individual neurons exhibited greater variance (Figure 7D) and the linear fits had a shallower slope than with widefield imaging (two-photon: − 0.63% deg^-1^, widefield: −1.04 % deg^-1^. No correlation was present between coherent motion response and azimuth (Figure S6). Taken together, the individual neurons largely reflect both the areal differences and retinotopic asymmetry observed widefield imaging, while also revealing a bias toward horizontal coherent motion that was not evident with mesoscale imaging.

**Figure 7:**
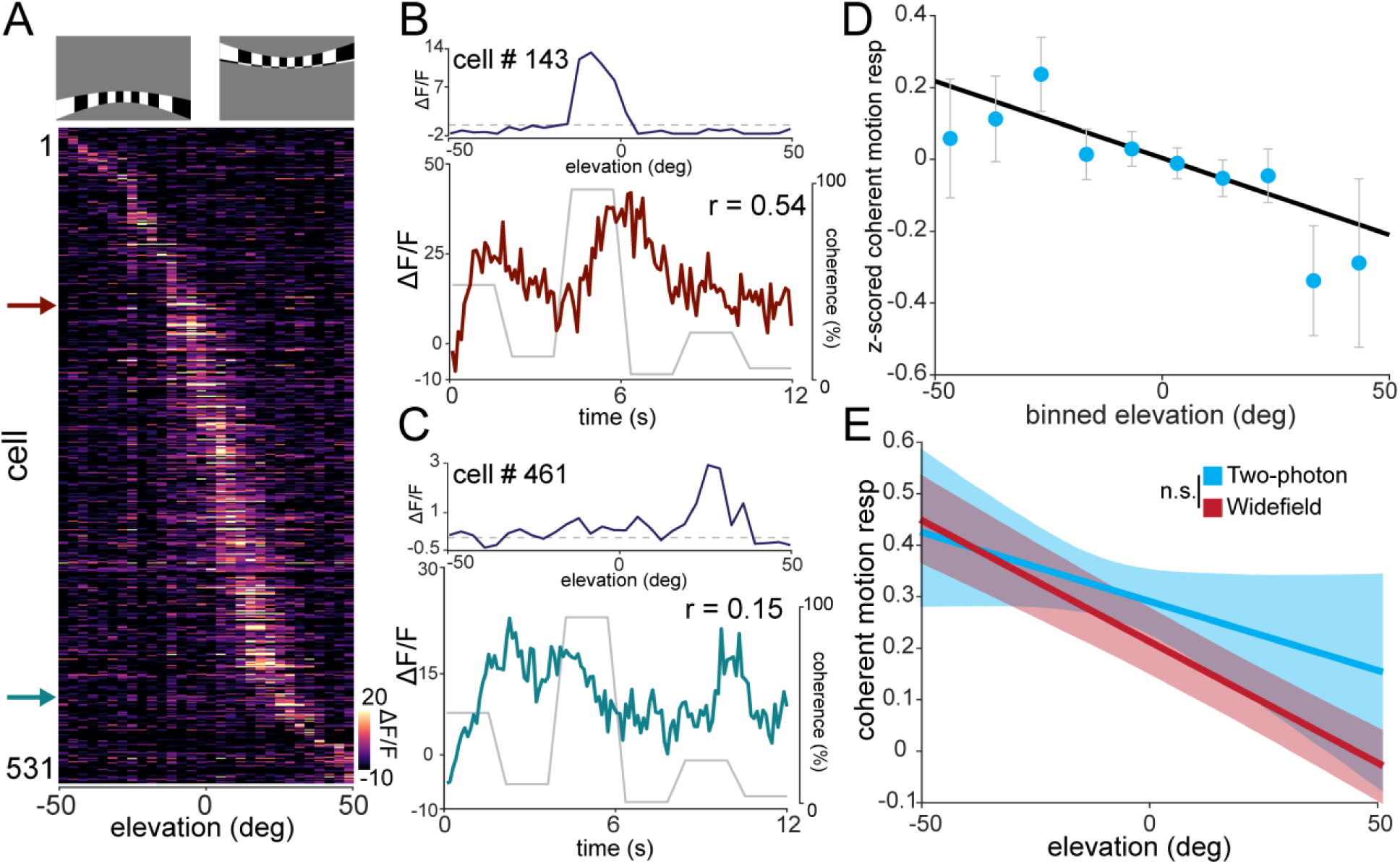
Retinotopic asymmetry of single cell coherent motion responses in V1. **(A)** Top: examples of a high elevation and low elevation spherically corrected retinotopic bar stimulus. Bottom: Averaged responses of each cell in the field to the elevation retinotopic bar stimulus, ordered by elevation preference. **(B and C)** Top: the preferred elevation location of the specified cell. Bottom: The calcium response of the cell during the RDK stimulus, with the coherence trace overlaid. **(D)** Plot of elevation preference versus z-scored coherent motion response with neurons averaged within 10**°** elevation bins. Error bars are mean ± s.e.m. **(E)** Plot of elevation preference versus coherent motion response averaged across experiments for widefield (red) and 2-photon experiments (blue). The confidence band represents the bootstrapped 95% confidence intervals of the slope and intercept.

## DISCUSSION

In this paper, we report three key findings: (1) all visual areas measured, including V1, exhibit reliable responses to global coherent motion independent of the spatial content of the retinal image, (2) HVAs respond heterogeneously to coherent visual motion, with stronger responses in putative dorsal stream areas, and (3) neurons which represent the inferior visual field having significantly stronger responses to coherent visual motion across visual areas. These findings have implications both for our understanding of murine visual motion processing and for more generalized principles of cortical organization.

### Role of V1 and HVAs in coding of coherent motion

In highly visual mammals, researchers have found defined extrastriate regions that are specialized for processing of coherent visual motion, including posteromedial (PMLS) and posterolateral (PLLS) lateral suprasylvian cortex in cats [49,50], MT and MST cortex in non-human primates [9,12–14], and V5 in humans [51]. Anatomical [40,43,44] and functional [23,45] evidence has suggested that mouse visual cortex might also have dedicated areas specialized for processing of coherent motion, but previous studies had not tested this possibility systematically.

In this paper, we used widefield calcium imaging to systematically investigate the responses of 7 visual areas (V1, LM, AL, PM, LI, RL, and AM) to coherent motion stimuli. We found that all visual regions exhibited reliable responses to coherent motion, but that areas AM, AL, and PM exhibited stronger responses to coherent motion than RL, V1, LM, and LI (Figure 3). These findings held true at the single cell level, as shown with 2-photon calcium imaging of selected populations (Figure 6). This is mostly consistent with prior categorization of these regions into putative dorsal (AL, PM, RL, AM) and ventral (LM, LI) streams [40,43–45]. One exception is area RL, which is generally considered a dorsal stream area, but in our study had moderate responses to coherent visual motion (in between ventral and dorsal stream areas) using RDKs, though not natural movies. A recent paper suggests that region RL might be preferentially involved in processing binocular disparity, with inferior responsive neurons particularly attuned to the near field visual stimuli rather than visual motion [52]. Another possibility is that the stimulus parameters used for the RDKs are not well suited to driving RL neurons, but that coherent motion found in natural movies is more effective.

Although we do not currently know how downstream cortical regions use signals from visual HVAs in local computations, the cortico-cortical connectivity of the motion responsive regions suggests an intriguing possibility. The HVAs with strong coherent motion responses (PM, AL, and AM) are highly interconnected with regions involved in navigation, including the parietal cortex, retrosplenial cortex, anterior cingulate cortex, and the presubiculum [44]. Indeed, area AM is often considered a posterior parietal region [43] though there is not yet widespread agreement on the boundaries between HVAs and parietal cortices [53]. This connectivity profile suggests that neurons in these HVAs may play an important role in visually-guided navigation, providing external motion cues to higher cortical regions.

### Retinotopic asymmetry in coding of coherent motion

Although we found that mice are similar to cats and primates in having specialized HVAs dedicated to coherent motion processing, we also found a new principal of organization *within* visual areas that had not been described in previous studies. Specifically, we found that motion processing in mouse visual cortex is asymmetric across elevation, but not azimuth, with neurons representing the inferior field exhibiting significantly stronger coherent motion responses both across visual cortex and within defined visual areas (Figures 5 and 7). The inferior field bias is somewhat surprising since it is known that overhead motion (such as looming stimuli) can cause strong behavioral reactions such as freezing and escape responses [54]. Indeed, there may even be specialized RGCs for local motion in the superior visual field [29]. However, recent studies indicate that behavioral responses to overhead motion appear to be principally mediated by a direct circuit from retina to superior colliculus to brain stem nuclei [55,56]. This raises the possibility that motion-responsive RGCs may project to separate thalamic or collicular targets depending on their retinotopy.

It has previously been found that V1 projections to different HVAs exhibit visual response properties consistent with the target area [57]. One way the dorsal stream areas (AL, PM, RL, AM) could inherit both an inferior bias in visual field coverage [40] and strong responses to coherent motion relative to other HVAs would be if they received preferential innervation from anterior V1; with ventral stream areas (LM, LI) receiving preferential input from posterior V1.

An open question is whether there is a functional reason for the inferior field bias in coherent motion responses. One possibility is that coherent motion responsive units convey external motion cues to downstream regions, which would be consistent with known cortico-cortical connectivity [44]. Since mice have coarse spatial resolution (∼10° receptive field size [58]) and their eye level is close to the ground, cues for external motion are likely more prevalent in the inferior visual field. That the horizontal bias in preferred direction observed in single neurons (Figure 6D-F) also reflects a bias in the inferior field external motion cues reaching the visual system is consistent with this interpretation.

These results also raise the question of whether the inferior field bias of coherent motion responses might also exist in other visual mammals such as primates. Since primates have higher acuity and their eye level is farther from the ground, it is possible that external motion cues are more distributed across the visual field than for mice, obviating the need for asymmetric processing of motion cues across retinotopic space. However, behavioral evidence of an “inferior field advantage” in detecting motion suggests that there may indeed be some inferior field bias present even in humans [59,60], though this has not been confirmed at the cellular level.

It should be noted that although coherent motion responses were stronger in particular HVAs and in retinotopic regions corresponding to the inferior visual field at the population level, there was substantial heterogeneity at the level of individual neurons (Figure 6 and 7), and that coherent motion responsive neurons could be found throughout the retinotopic gradient and visual areas.

### Implications for organizational principles of cortical processing

One of the most surprising findings of the current study is that there is as much variability in coherent motion responses within a visual area as across visual areas. Several recent studies have found similar mesoscale organization in other response properties in mouse visual cortex. For example, recent studies have found retinotopic asymmetry of color selectivity within area V1 [61] and binocular disparity preference within area RL [52]. Moreover, researchers also found that neurons in higher visual and parietal regions exhibit anatomical gradients of task-modulated responses during navigation in a virtual environment, and that these gradients cross retinotopic borders [62]. Together with our results, this suggests that visual input is not processed uniformly in each area as it progresses through the cortical hierarchy, but rather that there is considerable parallel processing along retinotopic axes *within* defined areas. Since topographic organization is present in many cortical areas, further research is necessary to determine whether this principle is present in other species and in other sensory, and non-sensory, cortices. These results suggest a general principle of sensory coding, which takes advantage of topographic mapping to selectively route visual information within areas throughout the cortical hierarchy, increasing the bandwidth for ethologically relevant stimuli across cortex.

## METHODS

### Animals

For cortex-wide calcium indicator expression, Emx1-Cre (Jax Stock #005628) × ROSA-LNL-tTA (Jax Stock #011008) × TITL-GCaMP6s (Jax Stock #024104) triple transgenic mice (*n* = 17) were bred to express GCaMP6s in cortical excitatory neurons. For widefield and 2-photon imaging experiments, 6 - 12 week old mice of both sexes (4 males and 13 females) were implanted with a head plate and cranial window and imaged starting 2 weeks after recovery from surgical procedures and up to 10 months after window implantation. The animals were housed on a 12 hr light/dark cycle in cages of up to 5 animals before the implants, and individually after the implants. All animal procedures were approved by the Institutional Animal Care and Use Committee at University of California, Santa Barbara.

### Surgical Procedures

All surgeries were conducted under isoflurane anesthesia (3.5% induction, 1.5 - 2.5% maintenance). Prior to incision, the scalp was infiltrated with lidocaine (5 mg kg-1, subcutaneous) for analgesia and meloxicam (1 mg kg-1, subcutaneous) was administered pre-operatively to reduce inflammation. Once anesthetized, the scalp overlying the dorsal skull was sanitized and removed. The periosteum was removed with a scalpel and the skull was abraded with a drill burr to improve adhesion of dental acrylic. A 4 mm craniotomy was made over the visual cortex (centered at 4.0 mm posterior, 2.5 mm lateral to Bregma), leaving the dura intact. A cranial window was implanted over the craniotomy and sealed first with silicon elastomer (Kwik-Sil, World Precision Instruments) then with dental acrylic (C&B-Metabond, Parkell) mixed with black ink to reduce light transmission. The cranial windows were made of two rounded pieces of coverglass (Warner Instruments) bonded with a UV-cured optical adhesive (Norland, NOA61). The bottom coverglass (4 mm) fit tightly inside the craniotomy while the top coverglass (5mm) was bonded to the skull using dental acrylic. A custom-designed stainless steel head plate (eMachineShop.com) was then affixed using dental acrylic. After surgery, mice were administered carprofen (5 mg kg-1, oral) every 24 hr for 3 days to reduce inflammation. The full specifications and designs for head plate and head fixation hardware can be found on our lab website (https://labs.mcdb.ucsb.edu/goard/michael/content/resources).

### Visual Stimuli

All visual stimuli were generated with a Windows PC using MATLAB and the Psychophysics toolbox [63]. Stimuli used for widefield visual stimulation were presented on an LCD monitor (43 × 24cm, 1600 × 900 pixels, 60Hz refresh rate) positioned 10 cm from the eye at a 30 degree angle to the right of the midline, spanning 130° (azimuth) by 100° (elevation) of visual space. For two-photon imaging of single cell responses, visual stimuli were presented on an LCD monitor (17.5 × 13 cm, 800 × 600 pixels, 60Hz refresh rate) positioned 5 cm from the eye at a 30 degree angle right of the midline, spanning 120° (azimuth) by 100° (elevation) of visual space.

Retinotopic mapping stimuli consisted of a drifting bar that was spherically corrected to account for visual field distortions due to the proximity of the monitor to the mouse’s eye [23]. A contrast-reversing checkerboard was presented within the bar to better drive neural activity (0.05 cycles degree^-1^ spatial frequency; 2 Hz temporal frequency). The bar smoothly drifted at 10.8 degrees sec^-1^ and had a width of 8° of visual field space for elevation and 9° of visual field space for azimuth. The stimulus was swept in the four cardinal directions: left to right, right to left, bottom to top, and top to bottom, repeated 20-60 times.

Natural movies were a set of 22 home cage movies recorded from a mice with head-mounted cameras provided by E. Froudarakis and A.S. Tolias [48]. Each movie had a duration of 10 sec and was presented at a frame rate of 30 Hz. For each experiment, 3 movies were randomly selected from a pool of all movies and presented for 20 repeats in random order.

Random dot kinematograms (RDKs) consisted of black dots presented on a 50% gray screen [64]. Each dot had a diameter of 2° of visual space, and the number of dots was adjusted so that the screen was 20% occupied by dots. Dots moved at a speed of 80° sec^-1^ in a randomly assigned direction (0° to 315°, in 45° increments), and had a lifetime of 60 frames. A subset of these dots (3%, 6%, 12%, 24%, 48%, or 96%) had their directions adjusted to the same direction to create coherent motion within the random dot motion. The coherent motion trace, which dictates the amount of coherence in the stimulus, was randomly generated at the beginning of the stimulus presentation and remained constant across repeats. Other stimulus features such as coherence direction and dot position were randomized across trials in order to eliminate reliable responses to all visual features except coherent motion. For widefield stimulation, a square wave coherence trace consisting of a pseudorandom presentation of all coherence values was created for each experiment, with motion direction changing across repeats. For two-photon stimulation, a short pseudorandom coherence trace was generated, with coherence changing smoothly between target coherence values, and the RDK stimulus was repeated multiple times in different motion directions to allow measurement of directional tuning. A 2 sec blank gray screen preceded each trial, and the stimulus was repeated for 10-20 trials.

To map azimuth and elevation preferences, full screen length bars (width = 20°) of a contrast-reversing checkerboard (spatial frequency of 0.04 cycles degree^-1^; temporal frequency = 5 Hz) were displayed on a 50% gray screen. There were 30 overlapping bar locations for horizontal bars (elevation mapping) and 40 overlapping bar locations for vertical bars (azimuth mapping). The bar appeared at each location in random order for 1 s, with a 2 s gray screen between repeats.

### Widefield Imaging

After >2 weeks of recovery from surgery, GCaMP6s fluorescence was imaged using a custom widefield epifluorescence microscope. The full specifications and parts list can be found on our lab website (https://labs.mcdb.ucsb.edu/goard/michael/content/resources). In brief, broad spectrum (400-700 nm) LED illumination (Thorlabs, MNWHL4) was band-passed at 469nm (Thorlabs, MF469-35), and reflected through a dichroic (Thorlabs, MD498) to the microscope objective (Olympus, MVPLAPO 2XC). Green fluorescence from the imaging window passed through the dichroic and a bandpass filter (Thorlabs, MF525-39) to a scientific CMOS (PCO-Tech, pco.edge 4.2). Images were acquired at 400 × 400 pixels with a field of view of 4.0 × 4.0 mm, leading to a pixel size of 0.01 mm pixel^-1^. A custom light blocker affixed to the head plate was used to prevent light from the visual stimulus monitor from entering the imaging path.

### Widefield Post-processing

Images were acquired with pco.edge camera control software and saved into multi-page TIF files. All subsequent image processing was performed in MATLAB (Mathworks). The ΔF/F for each individual pixel of each image frame was calculated as:

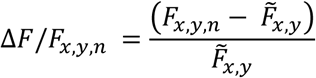

where *F*_*x,y,n*_ is the fluorescence of pixel *(x, y)* at frame *n*, and 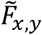 is defined as the median raw fluorescence value across the entire time series for pixel *(x, y)*. Subsequent analyses were performed on whole-frame ΔF/F matrices.

### Identifying HVAs using Widefield Retinotopic Mapping

For identification of HVAs, responses to drifting bar stimuli were averaged across each stimulus (horizontal left to right, horizontal right to left, vertical bottom to top, vertical top to bottom) [65]. Next, for each pixel, the phase of the first harmonic of the 1D Fourier transform was calculated to create retinotopic maps (phase maps) for each direction, which were then phase-wrapped to ensure smooth phase transitions between pixels. Lastly, to remove the delay due to the rise time of the GCaMP6s signal, phase maps of opposite directions (forward vs backwards, upward vs downward) were subtracted from one another [65].

Visual field sign maps were derived from the sine of the angle between the gradients in the azimuth and elevation phase maps. The resulting sign maps underwent post-processing as previously described [39,42]. Briefly, sign maps were first smoothed and thresholded, then each sign patch was dilated to fill in gaps between areas. Next, we applied an iterative splitting and merging process to further refine maps. First, each patch was checked for redundant coverage of visual space, and if significant redundancy (>10% shared visual field coverage) was found, the patch was split to create two separate patches. Conversely, adjacent same-sign patches were merged if they had little redundancy (<10% shared visual field coverage). After processing, borders were drawn around each patch, and resulting patches were compared against published sign maps for both size and sign to label each patch as a visual area. Visual areas V1, LM, AL, PM, LI, RL, and AM were present in all mice (Figure S1).

### Analysis of Natural Movie and RDK Stimuli

For natural movie reliability maps, we calculated the reliability of each pixel according to the following formula:

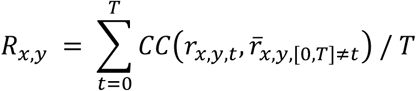

where *R* is reliability for pixel (*x, y*), *t* is the trial number from [0,*T*], *CC* is the Pearson correlation coefficient, *r*_*x,y,t*_ is the response of pixel *(x, y*) on trial *t*, and 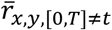 is the average response of pixel (*x, y*) on all trials excluding trial *t*.

For calculating coherent motion in natural movies, the pixel-wise motion vectors for each frame were calculated using the MATLAB optical flow toolbox (Mathworks). For each frame, the component vectors for each pixel were summed, and the magnitude of the summed component vectors was defined as the value of coherent motion for that frame.

In order to determine the uniformity of motion energy across the frame (Figure S4), we first calculated mean motion magnitude maps by taking the magnitude of each pixel’s motion vector for each frame, then meaning all resulting frames across a single movie. The uniformity of this image was gauged with a uniformity index, based on the ANSI standard for image uniformity. Briefly, the image is first divided into nine equal sections, which tile the image. The average brightness of each area is calculated. The uniformity index is then calculated as follows:

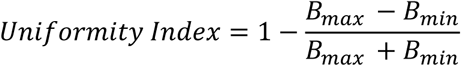

where *B* is the brightness of each section. Higher uniformity index values denote a more even image. As the “brightness” of each pixel in the mean magnitude map denotes the strength of motion information, a high uniformity index signifies even motion information across the screen.

For natural movies and RDKs, the coherent motion response of each pixel was calculated as the correlation of the mean response of the pixel and the coherent motion of the presented stimulus:

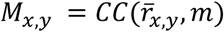

where *M*_*x,y*_ is the coherent motion response for pixel *(x, y), CC* is the Pearson’s correlation coefficient, 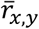 is the average adjusted ΔF/F response for pixel *(x, y)*, and *m* is the coherence value of the stimulus.

To combine reliability and coherent motion response maps across mice, individual maps were warped to align them to the Allen Brain Institute Common Coordinate Framework (https://scalablebrainatlas.incf.org/mouse/ABA_v3). Warping was performed on individual sign maps using custom code to ensure precise alignment of boundaries (available at: https://labs.mcdb.ucsb.edu/goard/michael/content/resources). The image transformation parameters were then applied to the reliability or coherent motion response maps to warp them to the aligned sign maps. Warped maps were used for visualization, but all statistical analyses were performed on individual maps, as described below.

Regions of interest for seven visual areas (V1, LM, AL, PM, LI, RL, and AM) were individually defined for each mouse using the mouse-specific sign map as previously described [40]. Pixels within each defined area were averaged to compare areal responses across all imaged mice.

The retinotopic dependence of coherent motion responses was calculated as the correlation between the retinotopic preference (elevation or azimuth) and the coherent motion response for all pixels within each area. Density plots were created using scatplot (MATLAB Central File Exchange). Briefly, a subsampled scatter plot was first created, then turned into a Voronoi diagram. The density of points within each Voronoi cell centered on the subsampled scatter plot was used to determine the density of the distribution in that portion of the scatter plot.

### 2-Photon imaging

After >2 weeks’ recovery from surgery, GCaMP6s fluorescence was imaged using a Prairie Investigator 2-photon microscopy system with a resonant galvo scanning module (Bruker). Prior to 2-photon imaging, epifluorescence imaging was used to identify the visual area being imaged by aligning to areal maps measured with widefield imaging.

For fluorescence excitation, we used a Ti:Sapphire laser (Mai-Tai eHP, Newport) with dispersion compensation (Deep See, Newport) tuned to λ = 920 nm. For collection, we used GaAsP photomultiplier tubes (Hamamatsu). To achieve a wide field of view, we used a 16x/0.8 NA microscope objective (Nikon) at 1x (850 × 850 μm) or 2x (425 × 425 μm) magnification. Laser power ranged from 40–75 mW at the sample depending on GCaMP6s expression levels. Photobleaching was minimal (<1% min^-1^) for all laser powers used. A custom stainless-steel light blocker (eMachineShop.com) was mounted to the head plate and interlocked with a tube around the objective to prevent light from the visual stimulus monitor from reaching the PMTs. During imaging experiments, the polypropylene tube supporting the mouse was suspended from the behavior platform with high tension springs (Small Parts) to reduce movement artifacts.

### 2-Photon Post-processing

Images were acquired using PrairieView acquisition software and converted into TIF files. All subsequent analyses were performed in MATLAB (Mathworks) using custom code (https://labs.mcdb.ucsb.edu/goard/michael/content/resources). First, images were corrected for X-Y movement by registration to a reference image (the pixel-wise mean of all frames) using 2-dimensional cross correlation.

To identify responsive neural somata, a pixel-wise activity map was calculated using a modified kurtosis measure. Neuron cell bodies were identified using local adaptive threshold and iterative segmentation. Automatically defined ROIs were then manually checked for proper segmentation in a graphical user interface (allowing comparison to raw fluorescence and activity map images). To ensure that the response of individual neurons was not due to local neuropil contamination of somatic signals, a corrected fluorescence measure was estimated according to:

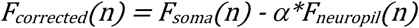

where *F*_*neuropil*_ was defined as the fluorescence in the region <30 μm from the ROI border (excluding other ROIs) for frame *n* and *α* was chosen from [0 1] to minimize the Pearson’s correlation coefficient between *F*_*corrected*_ and *F*_*neuropil*_. The ΔF/F for each neuron was then calculated as:

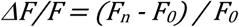

Where *F*_*n*_ is the corrected fluorescence (*F*_*corrected*_) for frame *n* and *F*_*0*_ defined as the mode of the corrected fluorescence density distribution across the entire time series.

### Analysis of 2-Photon Imaging Data

To map azimuth and elevation preferences, responses were measured during presentation of horizontal or vertical bars containing an alternating checkerboard stimulus. To identify neurons as visually responsive, a one-way ANOVA was first performed to screen for significantly preferential responses for specific stimulus locations. For neurons passing this criterion, we fit the responses with a 1D Gaussian model to determine the preferred azimuth and elevation. Finally, the receptive field preference of all significantly responding neurons in the imaging field were correlated to their pixel distance from the edge of the screen to ensure that the imaged neurons had receptive fields on the screen, and that the retinotopic preference correlated with the anatomical axis of azimuth/elevation (to ensure that the imaging field did not cross areal boundaries). Sessions that failed to exhibit a correlation between receptive field preference and pixel distance along the retinotopic axis were not used for further analysis. The cutoff for correlation was calculated for each recorded field by shuffling the cell locations 5000 times and calculating the correlation between receptive field preference and shuffled pixel distance to probe the underlying distribution. The 99^th^ percentile of the shuffled distribution was then chosen as the threshold correlation, and sessions whose calculated correlations were below this value were discarded

For analysis of single neurons to RDKs, we analyzed responses to different motion directions separately. We first only used the motion direction that produced the highest Pearson correlation coefficient between the neural response and the coherent motion signal, as responses to null directions were generally weak. However, to calculate the net preferred direction of the neuron, we treated the coherent motion responses to each direction as a vector, and calculated the vector sum of all vectors. The orientation of the resultant vector was defined as the preferred direction of that neuron.

For measuring coherent motion responses by area, we imaged fields of neurons at 2x magnification (425 μm × 425 μm) in identified visual areas, as identified in widefield visual field sign maps. We then compared distributions of single-cell coherent motion response across areas. For measuring the relationship of coherent motion responses to retinotopy in single neurons, we calculated the correlation coefficient between the coherent motion response and the preferred elevation.

### Statistical Information

To test statistical significance of single groups compared to a group of zero mean, single-sample t-tests were performed. For comparing experimental groups, two-sample paired and unpaired t-tests were performed for paired and unpaired groups, respectively. All t-tests were performed as two-tailed t-tests. To test for circular non-uniformity, we use the Hodges-Ajne test for angular uniformity.

## ACKNOWLEDGEMENTS

We would like to thank Emmanouil Froudarakis and Andreas S. Tolias for providing head-mounted natural movie videos and Ikuko T. Smith, Spencer L. Smith, Ming Hu, and Ben E. Reese for comments on manuscript. This work was supported by grants to M.J.G. from NIH (R00 MH104259), NSF (1707287), the Hillblom Foundation, and the Whitehall Foundation.

## AUTHOR CONTRIBUTIONS

K.K.S. and M.J.G. designed the experiments and analyses. K.K.S. conducted the experiments and analyzed the data. K.K.S. and M.J.G. wrote the manuscript.

## DATA AVAILABILITY

The data and all custom MATLAB analysis code that support the findings of this study are available from the corresponding authors upon request.

## SUPPLEMENTARY INFORMATION

### Supplemental Figures

**Figure S1.**
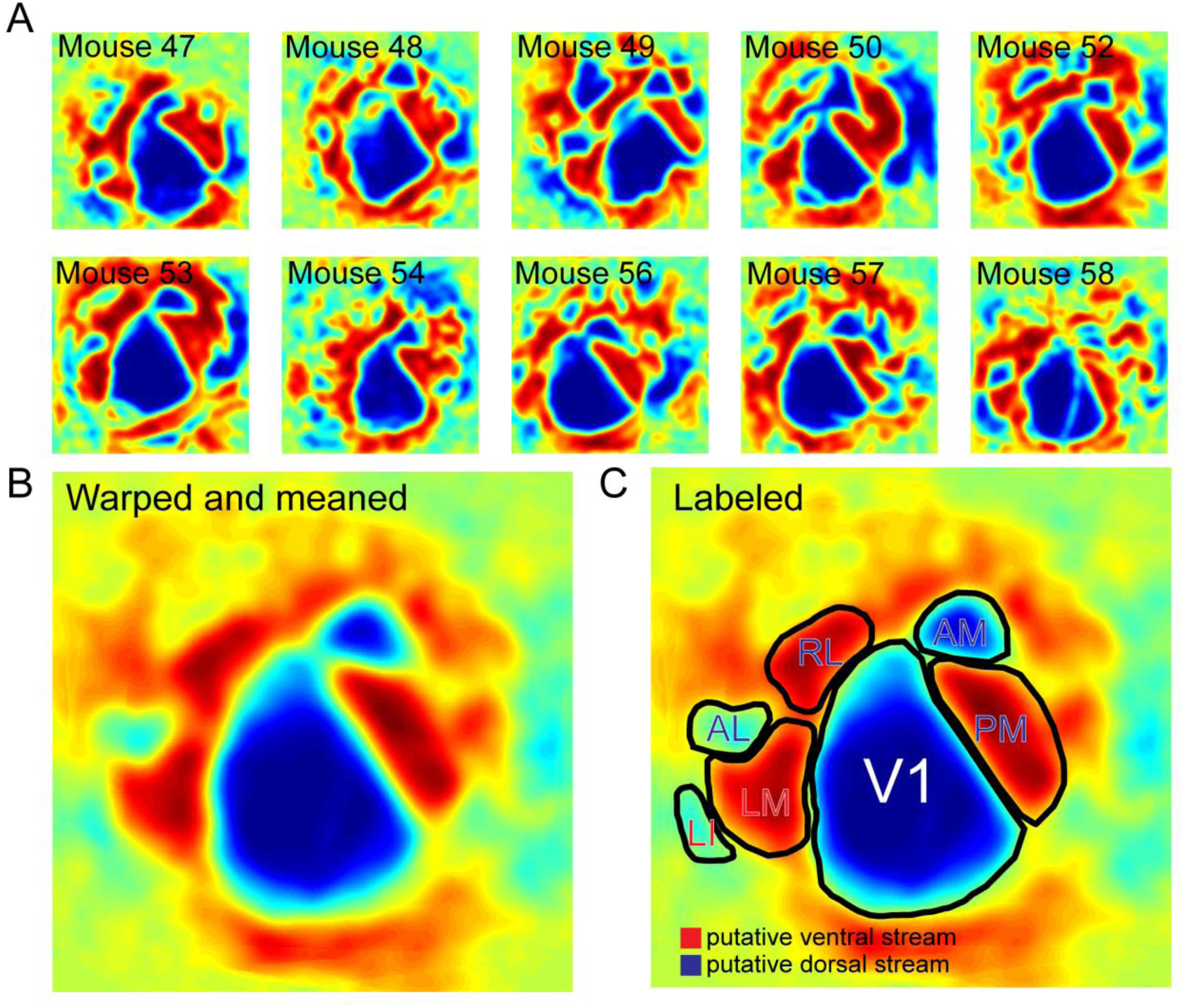
Related to Figure 1: Visual field sign maps from all imaged mice. **(A)** Sign maps for each mouse imaged for all ten mice used in both widefield RDK and natural scene experiments. All sign maps contain the seven areas analyzed (V1, LM, PM, AL, LI, RL, and AM). **(B)** Warped and averaged sign maps of all mice. **(C)** Same as in (B), but with areal boundaries and labels included. The putative mouse dorsal and ventral streams are designated in blue and red, respectively.

**Figure S2.**
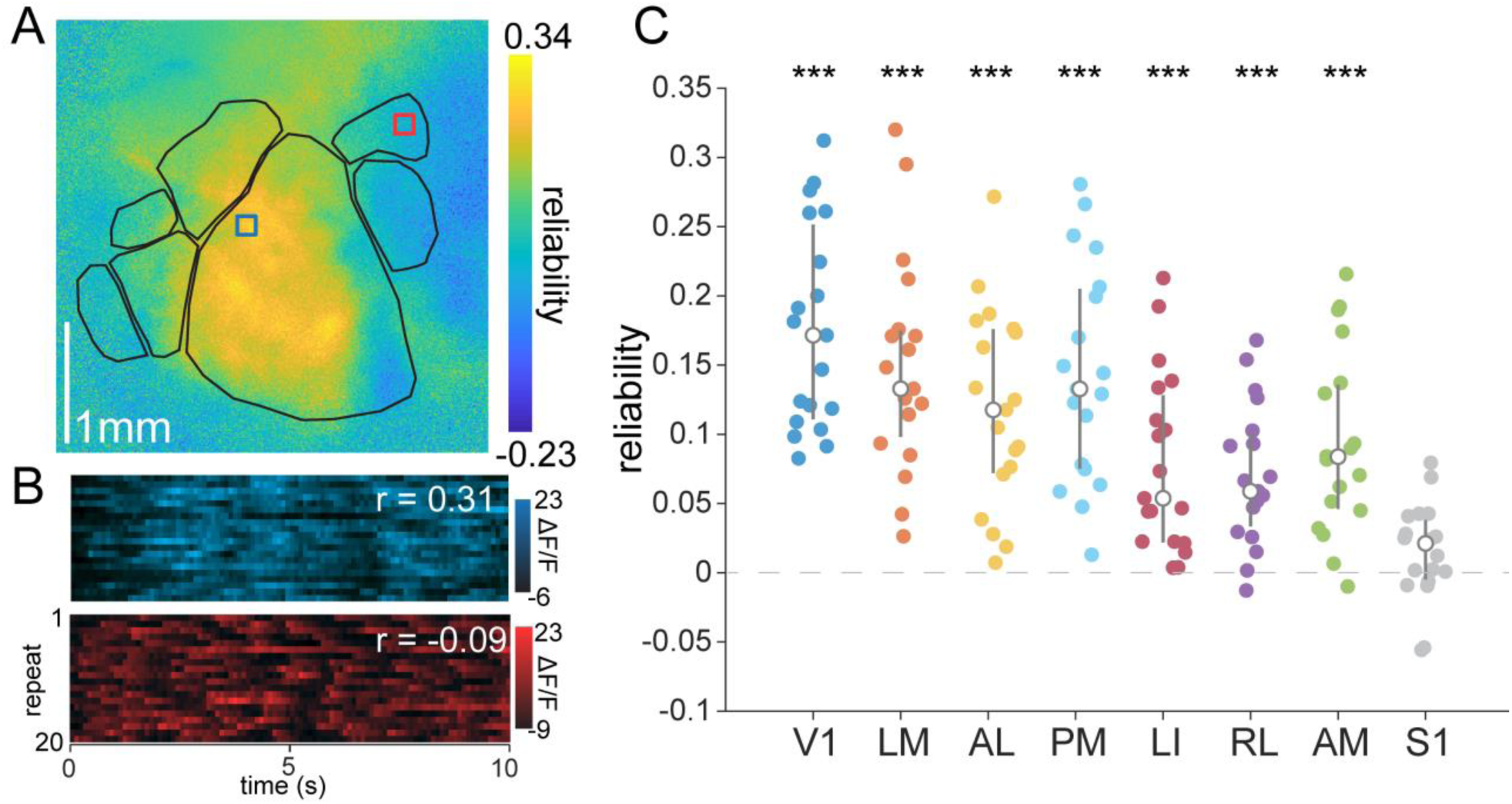
Related to Figure 1: Natural scene reliability across mouse visual cortex. **(A)** Map of reliability to natural scene stimuli. **(B)** Two example pixels from the labeled boxes in (A). The blue pixel shows from area V1 shows high reliability; whereas, the red pixel from area AM shows low reliability. **(C)** V1 and HVAs ordered in approximate ascending hierarchical order, showing a clear decrease in reliability. All areas but area S1 (have significantly reliable visual responses to the natural scenes (***: *p* < 0.001, V1: *p =* 4.92 ×10^−9^, LM: *p* = 1.08 × 10^−7^, PM: *p* = 2.50 × 10^−7^, AL: *p* = 9.14 × 10^−7^, LI: *p* = 4.44 × 10^−5^, RL: *p* = 9.31 × 10^−6^, AM: *p* = 7.47 × 10^−6^, S1: *p* = 0.079; single-sample t-test; *n* = 19 sessions over 7 mice).

**Figure S3.**
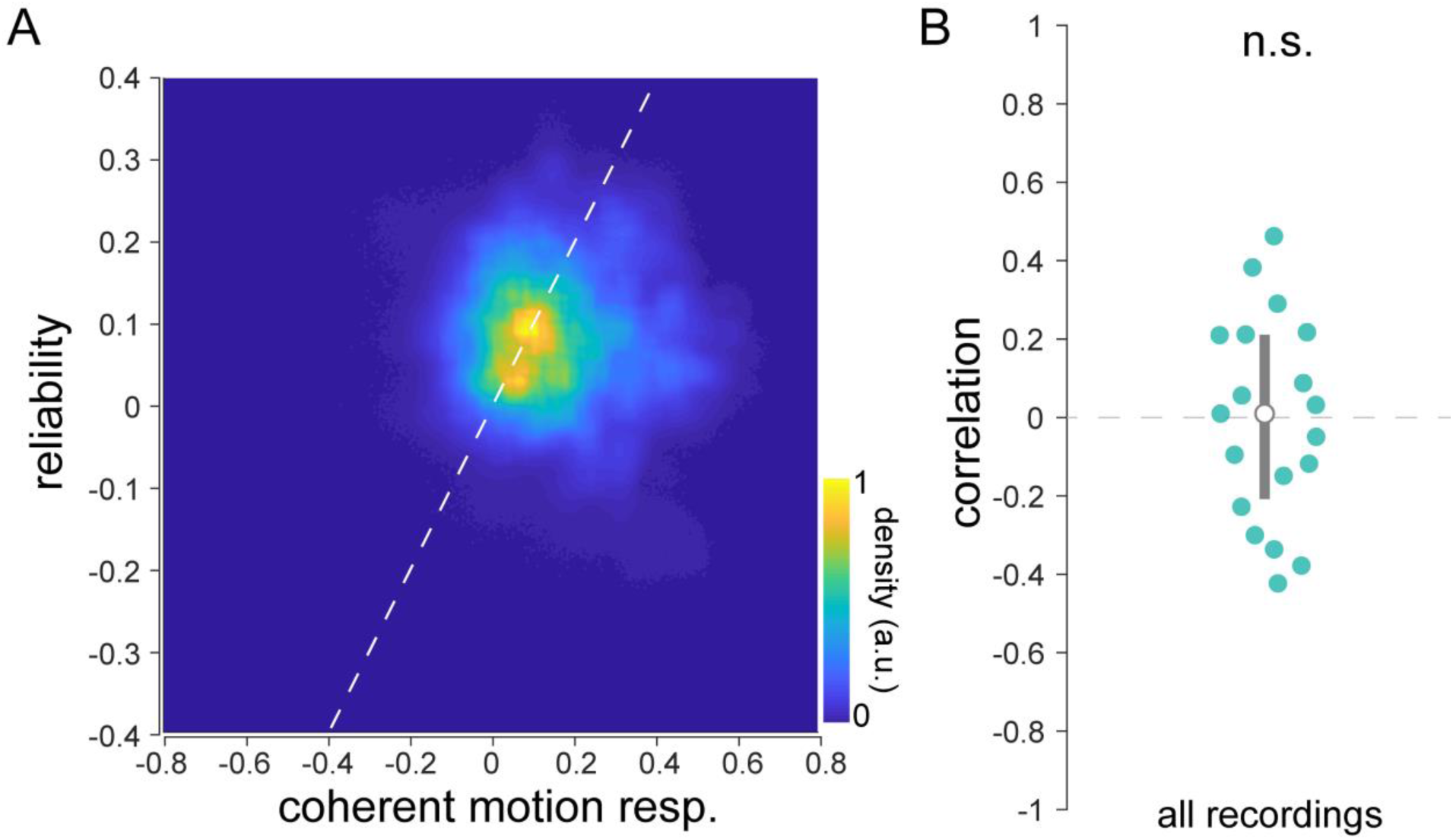
Related to Figure 1: Correlation of reliability and coherent motion response of natural scenes. **(A)** Density plot of reliability versus coherent motion responsiveness for all sessions. **(B)** Correlation between reliability and coherent motion response for each session (*n* = 19 sessions over 7 mice). There is no significant correlation between reliability and coherent motion response across sessions (*p* = 0.91; single-sample t-test).

**Figure S4.**
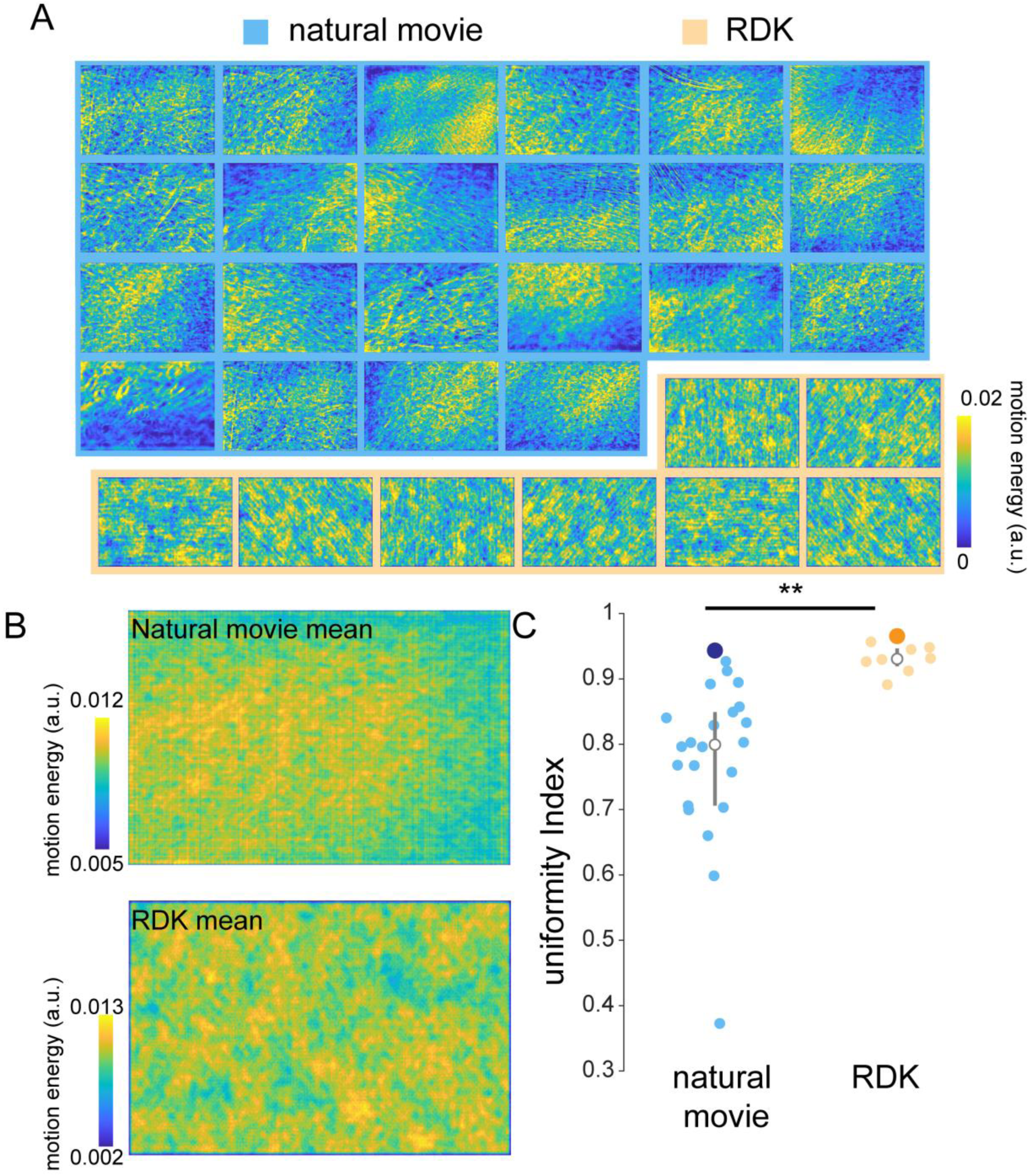
Related to Figure 1: Natural scenes have uneven motion energy across the frame. **(A)** Mean motion energy across all frames for each of 22 different natural scenes (blue background) and 8 RDKs of all shown directions (orange background). Whereas each natural scene has hotspots of motion energy, RDKs have motion energy present across the entire scene. When displayed across many directions, RDKs have an essentially flat distribution of motion energy across the frame. **(B)** Mean motion energy across all stimuli for natural scenes (top) and RDKs (bottom). In both cases, the combination of all stimuli results in a uniform distribution of motion energy across the frame. **(C)** Comparison of uniformity index (see experimental procedures) across the frame for the motion energy of natural scenes and RDKs. RDKs have much higher motion energy uniformity across the frame. For each stimulus, the large dot (dark blue for natural scenes, orange for RDKs) is the uniformity index of the single averaged motion energy from (B) (**: *p* < 0.01).

**Figure S5.**
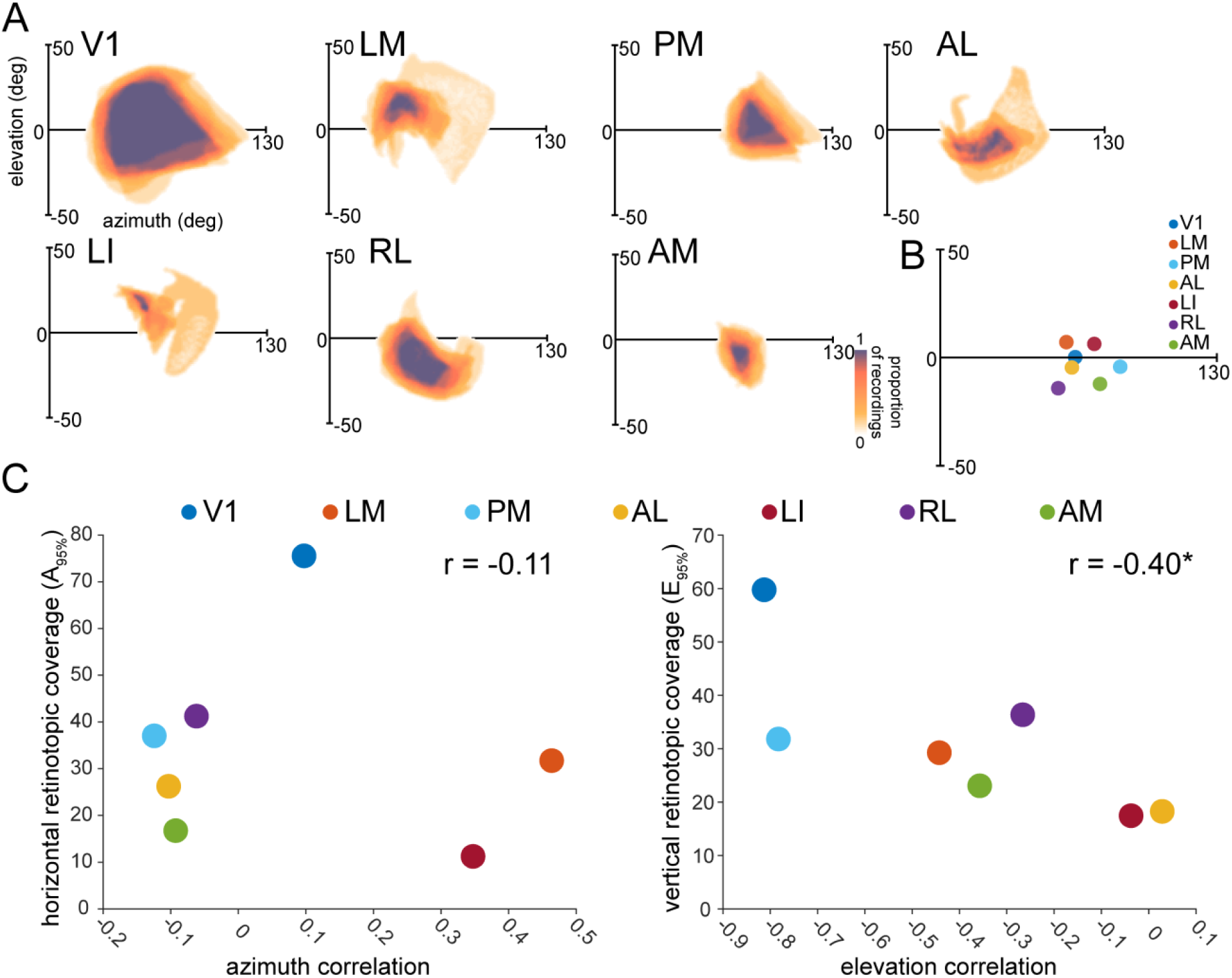
Related to Figure 5: Relationship between retinotopic correlation and visual coverage. **(A)** Visual field coverage of each identified visual area. The center of the axes is the center of V1, denoting (0, 0) in a monocular retinotopic space. **(B)** Centers of each identified area, showing the varying biases of each visual area to different parts of the visual field. **(C)** Comparison between the retinotopic correlation and retinotopic coverage (A_95%_) in its respective axis for azimuth (left) and altitude (right). Azimuth correlation does not depend on the visual span of each area in the azimuth; whereas, the altitude correlation decreases strongly with decreasing vertical span (azimuth: *p* = 0.81, altitude: *p* = 6.66 ×10^−4^; *n* = 10 sessions over mice, single-sample t-test).

**Figure S6.**
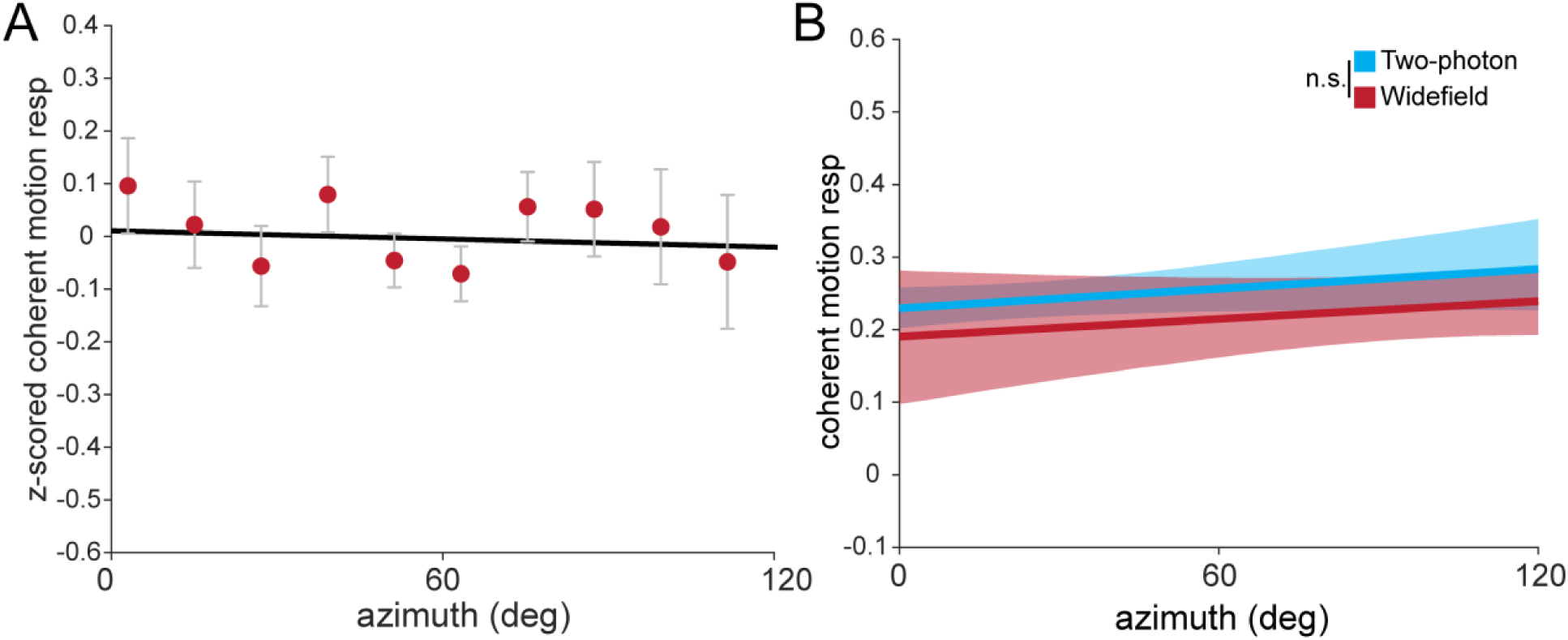
Related to Figure 7: Coherent motion responsiveness across azimuth in V1. **(A)** Plot of azimuth preference versus z-scored coherent motion response, averaged in 10**°** azimuth bins. Error bars are mean ± s.e.m. **(B)** Plot of azimuth preference versus coherent motion response averaged across experiments for widefield (red) and two-photon (blue) experiments, as in Figure 7E. Confidence band represents bootstrapped 95% confidence intervals of slope and intercept. The slopes between widefield and two-photon fit lines are not significantly different (*p* = 0.95, single-sample t-test).

**Supplemental Video S1: Widefield responses to retinotopic mapping stimuli**

Average widefield ΔF/F responses during presentation of horizontally or vertically drifting bars used in the retinotopic mapping procedure (scale bar indicates % ΔF/F). The white outlines indicate visual area boundaries based on the sign mapping procedure. Note the multiple waves of propagating activity resulting from discrete retinotopic maps. Inset: retinotopic mapping stimulus shown contralateral to the imaging window.

**Supplemental Video S2: Two-photon responses to RDKs**

**(A)** RDK stimulus shown to contralateral eye from two-photon session. **(B)** Two-photon imaging plane showing multiple cells responsive to the visual stimuli. Inset: Zoomed in view of a single neuron whose trace is highlighted above. **(C)** Comparison of the coherence signal from the RDK (top) with the neural activity of the highlighted neuron (bottom), showing the tight coupling of neural activity to coherent motion for this neuron.

